# Corticosterone release in very young siblicidal seabird chicks (*Rissa tridactyla*) is sensitive to environmental variability and responds rapidly and robustly to external challenges

**DOI:** 10.1101/2024.02.25.581949

**Authors:** Z.M. Benowitz-Fredericks, A.P. Will, S.N. Pete, S. Whelan, AS Kitaysky

## Abstract

In birds, patterns of development of the adrenocortical response to stressors vary among individuals, types of stressors, and species. Since there are benefits and costs of exposure to elevated glucocorticoids, this variation is presumably a product of selection such that animals modulate glucocorticoid secretion in contexts where doing so increases their fitness. In this study, we evaluated hypothalamic-pituitary-adrenal (HPA) activity in first-hatched free-living seabird nestlings that engage in intense sibling competition and facultative siblicide (black-legged kittiwakes, *Rissa tridactlya*). We sampled 5 day old chicks (of the ∼45 day nestling period), a critical early age when food availability drives establishment of important parent-offspring and intra-brood dynamics. We experimentally supplemented parents with food (“supplemented”) and measured chick baseline corticosterone secretion and capacity to rapidly increase corticosterone in response to an acute challenge (handling and a 15 min restraint in a bag). We also used topical administration of corticosterone to evaluate the ability of chicks to downregulate physiologically relevant corticosterone levels on a short (minutes) time scale. We found that 5 day old chicks are not hypo-responsive but release corticosterone in proportion to the magnitude of the challenge, showing differences in baseline between parental feeding treatments (supplemented vs non-supplemented), moderate increases in response to handling, and a larger response to restraint (comparable to adults) that also differed between chicks from supplemented and control nests. Topical application of exogenous corticosterone increased circulating levels nearly to restraint-induced levels and induced downregulation of HPA responsiveness to the acute challenge of handling. Parental supplemental feeding did not affect absorbance/clearance or negative feedback. Thus, while endogenous secretion of corticosterone in young chicks is sensitive to environmental context, other aspects of the HPA function, such as rapid negative feedback and/or the ability to clear acute elevations in corticosterone, are not. We conclude that 5 day old kittiwake chicks are capable of robust adrenocortical responses to novel challenges, and are sensitive to parental food availability, which may be transduced behaviorally, nutritionally, or via maternal effects. Questions remain about the function of such rapid, large acute stress-induced increases in corticosterone in very young chicks.

## 1. Introduction

The activity of the vertebrate hypothalamo-pituitary adrenal (HPA) axis is often considered in terms of costs and benefits. At a basal level, activity of the axis and the consequent release of glucocorticoid hormones is mandatory for survival, normal development (Wada, 2008), and modulation of metabolism (Jimeno and Verhulst, 2023), while the ability to rapidly increase glucocorticoids allows animals to maintain or reestablish homeostasis in the face of internal or external challenges (Sapolsky et al., 2000). While developmental exposure to glucocorticoids can increase fitness, presumably by better-preparing offspring for environmental conditions (Crino and Breuner, 2015), prolonged exposure to glucocorticoids during development (whether chronic or repeated acute elevations) can be associated with costs such as cognitive impairment, and reduced growth, immune function, and neural development (Kitaysky et al., 2003; Sapolsky and Meaney, 1986; Schoech et al., 2011). Thus, as with any process presumably shaped by selection, developing animals should only increase glucocorticoids when doing so increases fitness (i.e., benefits outweigh costs).

In birds, there is variation among species in the developmental stage at which chicks begin to respond to environmental challenges by activating the HPA axis and increasing circulating corticosterone (e.g. Adams et al., 2008; Bebus et al., 2020; Wada et al., 2007), as well as variation in which stimuli can elicit an HPA axis response from nestlings (e.g. Lynn and Kern, 2018, 2014). Many altricial and some semi-altricial species have a post-hatch “hypo-responsive” period (the “stress non-responsive period” in mammals) of relatively low HPA axis responsiveness (Bebus et al., 2020; Blas et al., 2006; Fiske et al., 2013, Jones et al., 2021; Sims and Holberton, 2000; Tilgar et al. 2009; Wada et al., 2009, 2007). This developmental pattern is thought to protect young chicks who are both vulnerable to excessive glucocorticoid exposure and too early in development to benefit from elevated glucocorticoids because they have life-histories that leave them unable to change physiology or behavior in sufficiently rapid or appropriate ways to increase fitness (Fiske et al., 2013; Sims and Holberton, 2000; TorresLMedina et al., 2019; Wada, 2008; Wada et al., 2007). However, some species have life-histories that involve complex early-life social dynamics that may be influenced by glucocorticoids (reviewed in Smiseth et al., 2011) and would therefore be expected to activate the HPA axis early in life.

For nest bound chicks that compete with siblings for limited parental resources via begging, aggression, and, in some species, siblicide, corticosterone may be a relevant mediator of social behavior (Loiseau et al., 2008; Vallarino et al., 2006). While glucocorticoids in adults are thought to help mediate the tradeoffs associated with allocating resources and behavior between self-maintenance and reproduction to maximize fitness (Bókony et al., 2009), chicks should be selected to maximize survival and optimize somatic growth such that they can successfully recruit to the breeding population. Food availability may be one of the most important factors that affects chicks’ growth and survival, behavior, and adrenocortical activity (Kitaysky et al., 2003). In some species, presumably those with few options available to increase food intake, food availability has minimal impacts on corticosterone release (Kitaysky et al., 2005). However, in other species, survival, and optimal growth in the face of limited food availability can evoke intense intra-brood competition at a very young age (reviewed in Morandi and Ferrer, 2014, Osorno and Drummond, 2003). These species may be selected for HPA activation early in postnatal life, despite potential costs of elevating glucocorticoids during development.

Food availability affects sibling competition in semi-precocial black-legged kittiwake (*Rissa tridactyla;* hereafter “kittiwake”) chicks (White et al., 2010). Kittiwakes reside in cliffside nests with 0 - 2 siblings and rely on parental food provisioning for ∼40-45 days before fledging (Hatch et al., 2020). Both parents forage at sea and deliver regurgitated meals to chicks multiple times a day. Chicks engage in facultative siblicide, where the first-hatched chick (hereafter “A-chick”) often out-competes the second-hatched chick (hereafter “B-chick”) and monopolizes parental food resources. A-chicks accomplish this by interrupting B-chick begging and feeding bouts, driving them out of the nest with repeated attacks, or killing them outright (Braun and Hunt, 1983; reviewed in Dickins, 2021). Most siblicide occurs during the first 10 days post-hatch, with chicks capable of intense aggression just a few days after hatching (Braun and Hunt, 1983). While the proximate cause of death is rarely clear (often the B-chick is simply missing from the nest), in 2022, hour-long observations at 28 nests documented aggression in 13 of 28 nests and found that only 5 of those 28 B-chicks survived to fledge (ZMBF unpubl.) While there are dozens of hypotheses about the evolution of brood reduction, most of them center on maximizing fitness when resources are limited (Morandini and Ferrer, 2015). Thus, compared to more “helpless” altricial nestlings, the complex behavioral repertoire of young kittiwake chicks is likely to be highly sensitive to energy balance and resource availability, and therefore may be modulated by glucocorticoids.

Black-legged kittiwake physiology has been studied intensively, including several investigations of HPA activity (both baseline and restraint-induced elevations of circulating corticosterone) in captive and free-living chicks (Table 1). These studies confirm that food availability is likely to be an important driver of variation in kittiwake chick HPA activity. Captive studies demonstrated clear consequences of food intake (both quality and quantity) on HPA activity of older chicks (29-31 days) subject to several weeks of experimental diets (Kitaysky et al., 1999), with low quality/quantity diet treatments dramatically increasing both baseline and restraint-induced corticosterone, but there was no change in baseline corticosterone with age in captive chicks fed ad libitum (Table 1). However, these captive chicks were raised in isolation, without any of the natural, complicated social interactions with parents and siblings which may influence HPA activity. The interannual and intercolony differences observed in free-living 12–15-day old chicks (Brewer et al., 2010, 2008) was hypothesized to have been driven by differences in food availability. While experimental food supplementation to free-living kittiwake nests at a different colony did not generate large differences in corticosterone in at least some breeding seasons (Schultner et al., 2013; Young et al., 2017), the lack of treatment effect was attributed to the high food availability naturally available to control (non-supplemented) adults in those years.

**Table 1.**
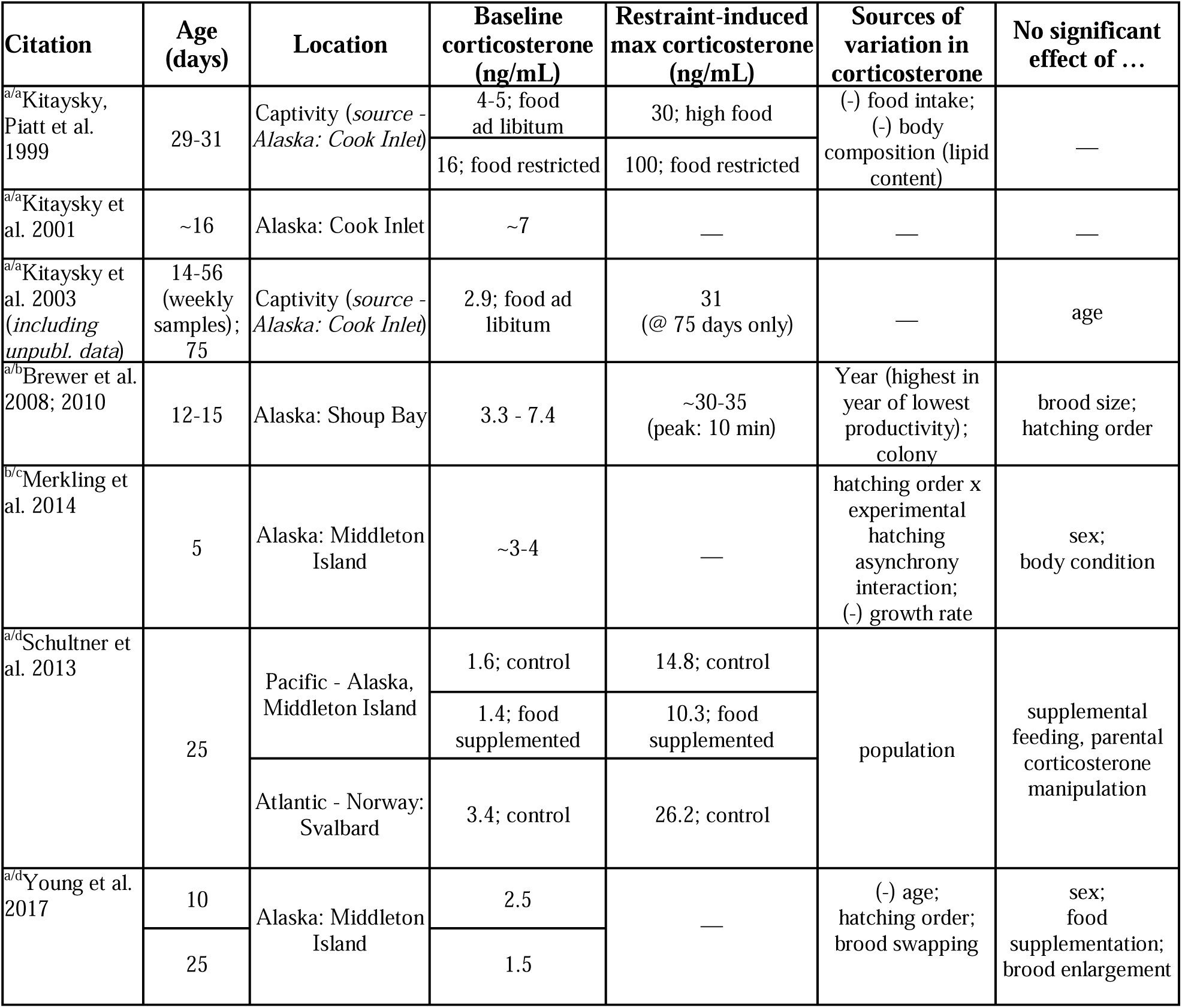
Review of studies measuring corticosterone in first-hatched black-legged kittiwake (*Rissa tridactyla)* chicks. Superscript letters indicate whether corticosterone radioimmunoassays were conducted using the same antibody (a = Esoterix, b = Biogenesis) and/or in the same lab (a = Wingfield, UW, b = O’Reilly, UP, c = Chastel, CEBC, d = Kitaysky, UAF). Note that comparison of absolute levels across studies may only be appropriate if hormone assays were conducted in the same lab using the same antibody, given the variation generated by inter-laboratory differences in extraction and assay methods, as well as specificity of antibodies used (Fanson et al. 2007). “∼” indicates that age is presented as approximate in some studies, or that hormone levels were estimated from figures.

Thus, overall, corticosterone release in kittiwake chicks appears to be responsive to environmental and social context at a variety of ages. However, all but one of these studies (Merkling et al., 2014; baseline corticosterone only) measured corticosterone in chicks that had already passed the most critical developmental window in terms of competition and resource monopolization, which occurs between age 3- and 10-days post hatch (Merkling et al. 2014). Intrabrood aggression drops to very low levels by 10 days post hatch (Leclaire et al., 2011; White et al. 2010), and 62-80% of B-chicks who died during the years of this study did so when both chicks were 10 days of age or younger (Whelan and Hatch, unpubl.). Thus, this early stage of development is critical for fitness, yet information about adrenocortical physiology at this age is lacking, a gap we sought to fill with this study. Because kittiwake life-history involves complex early-life social dynamics that may be influenced by glucocorticoids, we tested the hypothesis that the HPA axis of 5 day old black-legged kittiwakes is not “hypo-responsive” at very early developmental stages but is instead already sensitive to environmental context (interannual variability, experimental food supplementation) and physiological state (genetic sex, body condition). We measured baseline and restraint-induced corticosterone, as well as corticosterone uptake, negative feedback and clearance following exogenous corticosterone administration. We predicted that very young kittiwake chicks have well-developed adrenocortical activity, which is dampened by the experimental food supplementation of their parents.

## 2. Methods

### 2.1 Study site

This research was conducted in two years (2021 and 2022) at a free-living colony of black-legged kittiwakes nesting on a modified radar tower on Middleton Island, in the Gulf of Alaska (59°25’41.2“N 146°20’10.3”W). Details of the study site are described in Gill et al. (2002) and Gill and Hatch (2002), but in short: semi-partitioned nest sites on the outside face of the tower are each equipped with a sliding one-way mirrored glass window. The sliding glass provides access to individual nests from inside the tower for monitoring and manipulation with minimal disturbance to neighboring nests. All nests are monitored twice daily for events including egg-laying, hatching, chick death, and fledging. Chicks are not banded until day 25 post-hatch, so younger chicks are identified by hatching order using colored pen marks, applied on hatch day and refreshed every 5 days afterwards. Thus ages are known to the day. Tower nest sites are inaccessible to both terrestrial and aerial predators.

### 2.2 Food supplementation

The tower is also the site of a long-term food-supplementation experiment in which a subset of nests (hereafter “supplemented”) are provided with whole fish (either capelin (*Mallotus mallotus)* or herring *(Clupea spp.);* both preferred prey items) ad libitum, 3 times per day for the entirety of the breeding season (Gill et al. 2002), beginning several weeks before egg-laying. Adults readily consume the supplemental food; chicks consume the food supplement once they are large enough to consume whole fish (never observed prior to 7 days post-hatch, thus chicks in this study were not yet directly consuming the supplement). Adults at supplemented nests continue to forage at sea, but benefit from the supplementation in terms of reproductive success in most years (e.g. Kahane-Rapport et al., 2022). Occupants of control (hereafter “non-supplemented”) nests do not receive any supplemental food and rely solely on foraging at sea.

### 2.3 Blood sampling

In both years and regardless of treatment, all blood sampling occurred between 0600 and 0845, prior to the first daily supplemental feeding for supplemented nests at 0900. All chicks had been briefly handled only once before: on hatch day for measurements and marking. To prevent parents from startling abruptly and dislodging chicks, we started a timer then gently tapped on windows to alert parents and induce them to stand before retrieving a chick to take a blood sample. All blood samples were taken within 3 minutes of this initial tapping disturbance (2021: range = 38 - 180 sec, mean. = 82 sec, SD = 32 sec; in 2022: range = 45 - 172 sec, mean = 89 sec, SD = 25 sec). At each blood sampling event (hereafter “bleed”), the alar vein was punctured using a sterile 26G needle and 100-200 µL of whole blood was collected in heparinized microhematocrit tubes. Samples were refrigerated for up to 3 hours. They were then centrifuged to separate erythrocytes from plasma, both of which were frozen at -20°C for up to three weeks, then transported off the island and shipped on dry ice to the lab where they were stored at -80°C until use.

### 2.4 Experiment 1: Sources of variation in baseline corticosterone and response to restraint (Fig 1)

In 2021, between 24 Jun and 7 Jul, chicks from supplemented and non-supplemented nests containing two chicks were bled for baseline corticosterone (“undisturbed”; Table 2). Nests were sampled when the first-hatched A-chick was 5 days old (second-hatched B-chick ages ranged from 2 to 5 days old). For each nest, both A- and B- chicks were removed simultaneously and the A-chick was bled in < 3 min from initial disturbance. Chicks were then weighed and measured, B-chicks were returned to the nest and A-chicks were placed in a breathable cloth bag in proximity to a space heater, since 5 day old chicks do not have full thermoregulatory capacity. Fifteen min after initial disturbance, A-chicks were bled a second time to evaluate short-term adrenocortical response to restraint, then returned to their nest. Previous serial samples of restrained three-week old kittiwake chicks at this site took blood samples at < 3, 10, 30 and 50 min and found that corticosterone was highest at 10 min in some individuals, and at 30 min in others (Kitaysky, unpubl.). Because it is not possible to identify the timing of the true peak, we selected 15 min as a time point likely to capture variation in rates of increase, if not the peak increase. In 2022, baseline blood samples were collected from A-chicks between 29 June and 4 July for interannual comparisons.

**Figure 1.**
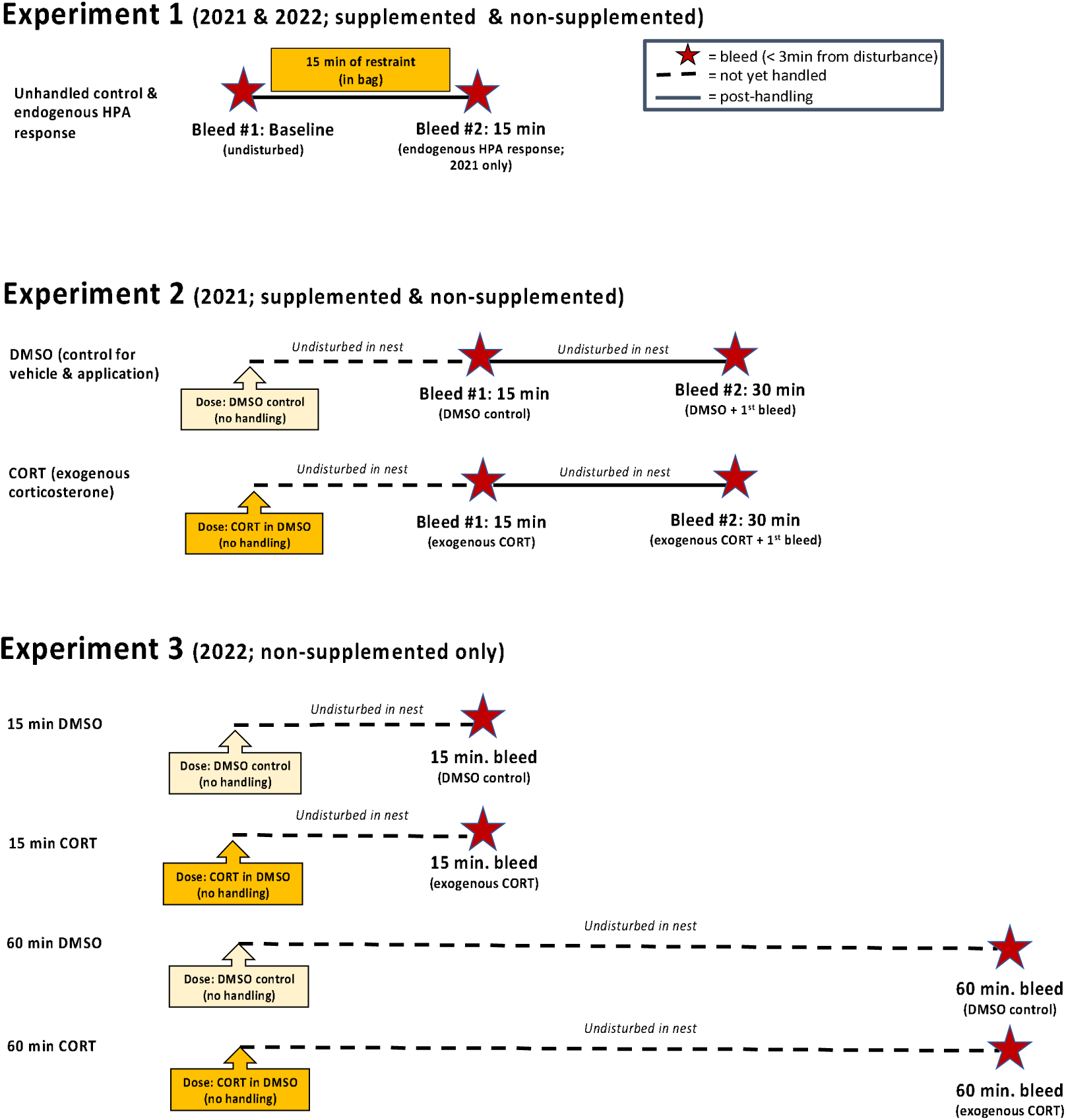
Experimental design. Timeline for blood sampling and topical hormone treatments of 5 day old black-legged kittiwake A-chicks.

**Table 2.** Circulating corticosterone levels (mean ± sd; ng/mL) in 5 day old, first-hatched black-legged kittiwake (*Rissa tridactyla)* y treatment and time. All chicks sampled on Middleton Island, Alaska. Parents at supplemented nests were provided with ish ad libitum 3x/day; non-supplemented relied only on natural food avilability. DMSO = treated topically with DMSO) and CORT = treated topically with corticosterone dissolved in DMSO. 15 min restraint values are from the same chicks baseline samples, and 30 min DMSO and CORT bleeds in 2021 are from the same chicks in the 15 min DMSO and CORT NA = no samples collected.

| Treatment/time | Year |  |  |  |
| --- | --- | --- | --- | --- |
|  | 2021 |  | 2022 |  |
|  | Parental food supplementation |  |  |  |
|  | Non- supplemented | Supplemented | Non- supplemented | Supplemented |
| Baseline (1st bleed) | 15.4 ± 10.9 (n=19) | 3.4 ± 1.9 (n=17) | 14.5 ± 15.6 (n=10) | 10.8 ± 7.5 (n=12) |
| 15 min restraint (2nd bleed) | 54.7 ± 10.7 (n=19) | 51.0 ± 17.9 (n=17) | NA | NA |
| 5 min DMSO (1st bleed) | 14.4 ± 12.6 (n=4) | NA | NA | NA |
| 5 min CORT (1st bleed) | 24.4 ± 20.5 (n=4) | NA | NA | NA |
| 15 min DMSO (1st bleed) | 11.0 ±7.7 (n=16) | 11.6 ±14.6 (n=11) | 13.7 ± 10.9 (n=11) | NA |
| 15 min CORT (1st bleed) | 41.0 ±22.8 (n=17) | 25.5 ± 19.2 (n=9) | 34.7 ± 16.7 (n=11) | NA |
| 30 min DMSO (2nd bleed) | 31.8 ± 17.0 (n=16) | 23.8 ±13.5 (n=11) | NA | NA |
| 30 min CORT (2nd bleed) | 57.2 ±20.9 (n=17) | 45.1 ±20.4 (n=9) | NA | NA |
| 60 min DMSO (1st bleed) | NA | NA | 6.4 ± 3.8 (n=14) | NA |
| 60 min CORT (1st bleed) | NA | NA | 27.7 ± 16.1 (n=14) | NA |

### 2.5 Experiments 2 and 3: Responses to exogenous corticosterone

To bypass some of the challenges and constraints associated with the use of subcutaneous corticosterone implants (Torres-Medina et al, 2018) and minimize changes in HPA activity associated with handling, we used topical administration of corticosterone in dimethyl sulfoxide (DMSO) to rapidly elevate corticosterone with minimal disturbance. This manipulation allowed us to evaluate differences in glucocorticoid uptake and/or clearance, as well as negative feedback. DMSO is a polar solvent that has been used in veterinary and human medicine as a delivery vehicle for topical application of lipophilic compounds for decades (Kumar and Darreh-Shori, 2017; Patil, 2013). It is considered non-toxic, though it can have off-target cellular effects such as inhibition of acetylcholinesterase, and is used by itself for a variety of conditions to reduce inflammation (Santos et al., 2003). In avian endocrinology, it has been used successfully for topical application of corticosterone (Busch et al., 2008; Pegan et al., 2019; Vitousek et al., 2018).

#### 2.5.1 Preparation of exogenous corticosterone treatments

Topical treatments were made following Pegan et al. (2019), though with a higher ratio of liquid DMSO to facilitate pipetting and surface area of contact on skin. In short, for the “CORT’’ treatment, 0.05 g corticosterone powder (Sigma #27840) was dissolved in 4 mL liquid DMSO, which was mixed with 6 mL veterinary grade DMSO gel to achieve a corticosterone concentration of 5 μg/μL. A control DMSO gel solution was made using the same ratio of liquid to gel DMSO, but with no corticosterone.

We used average body masses from 5 day old chicks measured in 2019 (90g) to anticipate that a 20 μL dose would deliver approximately 1 microgram of corticosterone per gram of body mass. This dose was selected based on previous experiments with topical administration of corticosterone in DMSO in birds (Busch et al, 2008, Pegan et al. 2019) in which it elevated corticosterone within a naturally occuring range. We used food coloring to visually confirm that vortexing rapidly and thoroughly dispersed liquid DMSO throughout the gel (though did not include dye in the final solutions). In the years of our study, average body mass of 5 day old chicks was lower than in 2019 (73g in 2021; 75g in 2022), such that 20μL of gel would have been a higher dose than intended. However, while 20μL of gel was drawn into each pipette tip, the viscosity of the gel meant that several drops were always left behind on the inside of the pipette tip. Repeated measurements in the lab using a microbalance to weigh pipetted DMSO gel mix (n = 20; compared to the weight of 20μL dispensed using a displacement syringe) indicated that an average of 12.4 ± 0.0019 µL of gel was delivered to each chick (57% of the dose originally calculated for 90g chicks). Thus, the average dose delivered to a 74g chick was 0.62 μg corticosterone/g body mass. We confirmed that this dose elevated corticosterone in kittiwake chicks to levels comparable to an endogenous response to handling in this study (see Fig. 4B)

#### 2.5.2 Administration of topical treatments

In 2021 and 2022, 5-day old - chicks from either one- or two-chick nests were treated with a drop of either corticosterone in DMSO gel (hereafter “CORT” treatment) or DMSO gel (hereafter “DMSO’’ treatment; see section 2.5.1). CORT and DMSO gels were kept in physically different locations (and away from where chicks were bled) with dedicated pipettes, tip boxes, and glove boxes, to minimize the risk of corticosterone contamination. Chicks were treated after tapping on the nest window to alert the parent and opening the window part way (causing the brooding parent to stand or briefly flush). We then reached into the nest with a pipettor set to 20 µL, and gently touched the tip to the skin on the back of the A-chick between the scapulae and behind the neck, dispensing the gel there (see supplemental material for video footage). We administered the gel to this location, where it is tucked behind the head when a chick is sitting, to minimize the likelihood that the gel was dislodged or transferred to brooding parents.

Application of the treatment usually took ∼15 seconds from initial disturbance of the nest site to completion (closing of window), unless the brooding parent was particularly aggressive, in which case it could take up to 30 seconds to work around the adult. Parents resumed brooding within seconds of window closure. Chicks were not weighed until the final blood sample was taken; chicks were excluded from further analyses *a priori* if they weighed < 50 g, a threshold below which chicks have dramatically reduced survival.

### 2.6 Experiment 2: Effects of exogenous corticosterone on endogenous production (Fig. 1)

We used A-chicks from food supplemented and non-supplemented nests containing one or two chicks (Table 2). We applied the topical gel treatment, took a blood sample 15 minutes after gel application and immediately returned the chick to the nest. Fifteen min after the first sample (i.e. 30 min after gel treatment), we removed the chick to take a second blood sample to evaluate the effect of exogenous corticosterone on subsequent endogenous corticosterone production in response to the handling that occurred during the first bleed. Finally, though DMSO is estimated to take at least 5 minutes to penetrate the skin and enter the bloodstream (Patil 2013), nothing is known about the acute time course of corticosterone uptake from topical DMSO application in young seabird chicks. Therefore, we checked for very rapid changes in corticosterone by treating a small subset of chicks with topical CORT or DMSO and taking a single blood sample 5 minutes after treatment administration (Table 2).

### 2.7 Experiment 3: Time course of exogenous corticosterone (Fig. 1)

In 2022, we repeated the topical CORT experiment (mixing fresh batches of gel solutions using the same protocol), but only used chicks from non-supplemented nests containing two chicks. Chicks were distributed across 4 treatment groups. Half of the topical CORT-treated chicks and half of the topical DMSO-treated chicks were bled for the first time 15 min after topical application to confirm rapid increase in corticosterone in CORT group, and the other half of the topically treated chicks were bled for the first time 60 min after application to evaluate chicks’ ability to clear exogenous corticosterone from circulation. Corticosterone levels in these four groups were compared to baseline levels in the same year (see Experiment 1, section 2.4).

### 2.8 Genetic sexing

All chicks were genetically sexed from DNA extracted from erythrocytes (Qiagen DNeasy Blood and Tissue kits, Cat # 69556). We used primers and protocols for PCR and gel electrophoresis described in Merkling et al. (2012) to differentiate the sexes by number of bands.

### 2.9 Corticosterone assays

Corticosterone was measured blind to treatment using radioimmunoassay, following the protocol described in Kitaysky et al. (2007). In short, plasma was equilibrated with tritiated corticosterone, extracted using dichloromethane, and assayed in duplicate. 2021 and 2022 samples were run in separate assays (three and one assays in 2021 and 2022, respectively), but all samples from the same chick were run in the same assay, with treatments distributed across assays. Each assay included a standard treated identically to samples to calculate interassay variability (interassay CV = 2.7%). Final corticosterone values were corrected for the percent of tritiated corticosterone lost during extraction (average recovery for 2021 = 96% +/- 3.18%, for 2022 = 92% +/- 2.96%).

### 2.10 Data analysis

To meet assumptions for parametric statistical tests, all corticosterone values were log-transformed, which normalized their distribution. Corticosterone concentrations among food, and CORT treatment groups had equal variance in all but one case (see below for modeling approach). We first determined whether brood size (one or two chicks) affected baseline corticosterone concentrations using a Welch’s two-sample t-test and tested for differences in body condition between years and feeding treatments (ANOVA) to account for potential confounding effects among explanatory variables on the measured physiological response (i.e. corticosterone concentration).

#### 2.10.1 Experiment 1: Sources of variation in baseline corticosterone and response to restraint

To test which ecological and biological factors affect 5-day old chicks’ baseline corticosterone and adrenocortical response to 15 min of handling and restraint (including whether there is a response) we used a model selection approach. We built an *a priori* candidate models of interest (Table 3) using a small set of variables that have been previously shown to affect adrenocortical response in birds: body condition, sex, feeding treatment, and in baseline adrenocortical function. We used Akaike’s Information Criterion (Burnham and Anderson, 2004) to identify the top performing model(s), defined as all models with AIC scores with a difference of < 2 from the top performing model. We calculated the effect size for the variables from each best fit model and considered any variable with an effect size whose 95% confidence interval does not overlap zero as statistically significant. We calculated AIC scores in R (v. 4.2.1, R Core Team 2022) using packages *AICcmodavg* (Mazarolle 2023). We calculated effect size using the package *effectsize* (Ben-Shachar et al., 2020), and tested variables in the top models for multicollinearity using the vif function in the *multcomp* package (Hothorn et al., 2008).

**Table 3.** Model selection results of variables affecting baseline corticosterone secretion of 5-day old chicks and *a priori* justification for model inclusion. Treatment = food supplementation to parents (food supplemented or non-supplemented). Top performing models are italicized.

| Model | K | AICc | Delta AICc | AICc Wt | *a priori* justification |
| --- | --- | --- | --- | --- | --- |
| *Treatment\*Year* | 5 | 38.93 | 0 | 0.56 | *Effects of parental nutritional status depend on interannual environmental variability* |
| *Year\*Treatment\*Body Condition* | 9 | 39.66 | 0.73 | 0.39 | *The effect of parental nutritional status on the relationship between chick energy stores and corticosterone varies interannually* |
| Full Two-Way Interaction Model* | 12 | 44.31 | 5.38 | 0.04 |  |
| Year+Treatment*Body Condition | 6 | 48.37 | 9.45 | 0 | Interannual environmental variation affects corticosterone, but parental nutritional status also modulates the impact of chick energy stores on corticosterone |
| Treatment | 3 | 49.35 | 10.43 | 0 | Parental nutritional status affects corticosterone |
| Treatment*Body Condition | 5 | 49.36 | 10.44 | 0 | Parental nutritional status modulates the impact of chick energy stores on corticosterone |
| Year+Treatment+Body Condition | 5 | 49.83 | 10.91 | 0 | Corticosterone is affected by interannual environmental variability, parental nutritional status, and chick energy stores |
| Treatment+Sex | 4 | 50.25 | 11.32 | 0 | Corticosterone is affected by parental nutritional status and chick sex. |
| Treatment+Body Condition | 4 | 51.27 | 12.35 | 0 | Both chick energy balance and parental nutritional status affect chick corticosterone |
| Year*Treatment+Body Condition+Sex | 6 | 51.36 | 12.44 | 0 | Corticosterone is affected by interannual environmental variability, parental nutritional status, chick energy balance and sex |
| Treatment*Sex | 5 | 52.39 | 13.46 | 0 | Parental nutritional status affects corticosterone in male and female chicks differently |
| Null (intercept only) Model | 2 | 62.82 | 23.9 | 0 | None of these variables explain variation in corticosterone |
| Year | 3 | 63.14 | 24.21 | 0 | Corticosterone reflects interannual environmental variability |
| Year+Body Condition | 4 | 63.5 | 24.58 | 0 | Corticosterone reflects both energy balance and interannual environmental variability |
| Body Condition | 3 | 63.82 | 24.9 | 0 | Corticosterone reflects energy balance |
| Year*Body Condition | 5 | 64.08 | 25.16 | 0 | The effect of chick energy balance on corticosterone is modulated by interannual environmental variability |
| Sex | 3 | 64.1 | 25.18 | 0 | Corticosterone reflects sex-specific differences |
| Body Condition*Sex | 5 | 64.14 | 25.22 | 0 | Corticosterone reflects energy balance in sex-specific ways |
| Year+Sex | 4 | 64.55 | 25.63 | 0 | Corticosterone reflects both sex and interannual environmental variability |
| Body Condition+Sex | 4 | 65.55 | 26.62 | 0 | Corticosterone reflects both energy balance and sex |
| Year*Sex | 5 | 66.76 | 27.83 | 0 | Male and female chicks respond differently to impacts of interannual environmental variability on corticosterone |

\*BCI+Sex+Treatment+Year+BCI\*Sex+BCI\*Treatment+BCI\*Year+Sex\*Treatment+Sex\*Year+Treatment\*Year

#### 2.10.2 Experiment 2: Effects of exogenous corticosterone on endogenous production

To test the effects of topical application of corticosterone via DMSO on chick’s subsequent adrenocortical response we ran a single mixed effects ANOVA testing whether corticosterone concentrations (both from the first and second bleed) differed between treatment groups. We included body condition as a covariate, and individual chick as a random effect. The variance in corticosterone levels at the second bleed was not equal among treatments, so we also accounted for unequal variance in the model. To test for the effect of feeding treatment on the response of chicks we ran two ANCOVA, one for corticosterone at the first bleed, and the other for corticosterone at the second bleed. For both models, feeding treatment was the predictor and body condition was a covariate.

#### 2.10.3 Experiment 3: Time course of exogenous corticosterone

We used the same mixed effects ANOVA model described above to test for differences in response over time to the exogenous treatments. To test for differences in the magnitude of the response between the first and second bleed in treatment groups we ran two difference ANCOVAs with the change in corticosterone concentration (i.e. delta corticosterone) as the response, corticosterone treatment (CORT or DMSO) and feeding treatment (supplemented or non-supplemented) as predictors and body condition as a covariate. The first was to examine change in corticosterone at comparable time intervals among the two exogenous treatments only. The second model included the delta corticosterone values for the control group (the difference between baseline and post-bag restraint). To test for differences in clearance of corticosterone over time among the treatment groups, for non-supplemented chicks only, we ran an ANCOVA with corticosterone concentration as the response, exogenous treatment as a predictor, and body condition as a covariate.

#### 2.10.4 Post-hoc analyses

We followed up all ANOVA and ANCOVA models with post-hoc comparisons and report p-values and F statistics from a Type III ANOVA using Satterthwaite’s method for modeled variables, and p-values from post-hoc Tukey contrasts. We analyzed data in R (v. 4.2.1, R Core Team 2022) packages *nlme* (for mixed effects ANOVA, Pinheiro and Bates 2023) and *multcomp* (for ANCOVA and post-hoc contrasts, Hothorn et al., 2008). We considered variables and contrasts with p-values < 0.05 as significant.

## 3. Results

### 3.1 Experiment 1: Sources of variation in HPA activity

#### 3.1.1 Sources of variation in baseline corticosterone

There was no difference in A-chick baseline corticosterone between one- and two-chick nests (t_18.16_= -0.06, p = 0.95), so brood size was omitted from subsequent analyses. Chick body condition did not differ between years (F_1,55_ = 1.69, p = 0.2), or feeding treatments (F_1,54_ = 1.77, p = 0.28). Model comparison indicated strong support for an effect of feeding treatment that differed between years, with both top models including the year*treatment interaction (Table 3). Post-hoc analyses of the top model indicated significant effects of year, treatment and their interaction - baseline corticosterone was higher in non-supplemented nests (effect size ± 95% confidence interval from top model: 0.99 ± 0.58, 1.41) but more so in 2021 than 2022 (-0.73 ± -1.16, -0.31), and was higher in chicks from both supplemented and non-supplemented nests in 2022 (0.57 ± 0.27, 0.87; Table 2, 3). The next best performing model included body condition; however, it was not an informative predictor on its own (-0.0042 ± -0.32, 0.31). Body condition did explain some of the variability with respect to feeding treatment (Fig 2) and so was included as a covariate in subsequent analyses.

**Figure 2.**
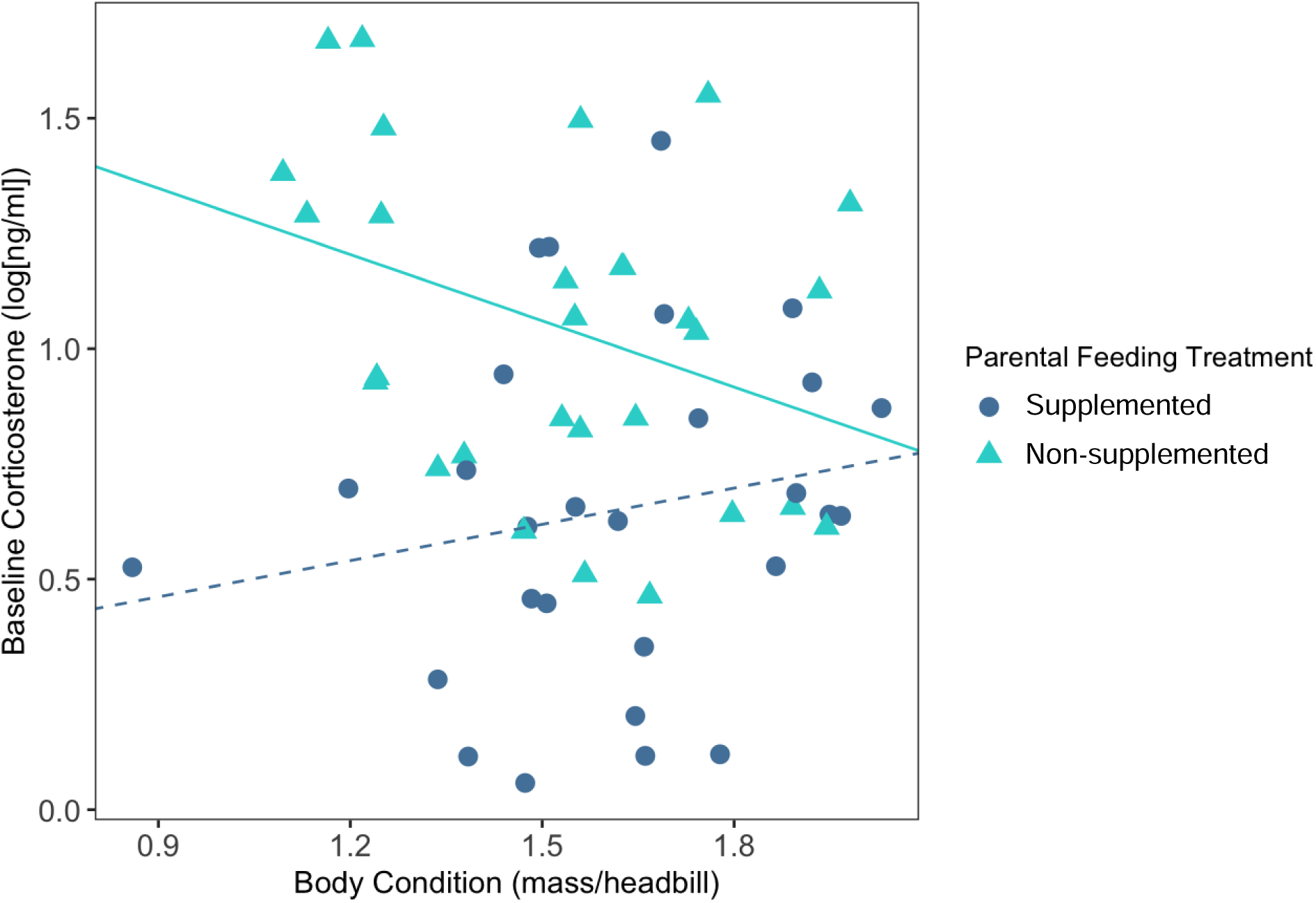
Body condition and baseline corticosterone. Body condition of 5 day old first-hatched kittiwake chicks was negatively related to baseline corticosterone levels, but this relationship was eliminated in nests where parents were provided with supplemental food (“supplemented”). Solid line = significant, dashed = non-significant.

#### 3.1.2 Endogenous corticosterone secretion in response to restraint

After 15 minutes of handling and restraint in a bag, 5-day old chicks increased corticosterone substantially (Table 2, Fig. 3, 4). The top model for corticosterone after 15 min of restraint included the interaction between baseline corticosterone and feeding treatment (effect size: 1.55 ± 0.65, 2.44; Table 4). In nests where parents were supplementally fed, higher baseline corticosterone was associated with lower HPA response - this was not the case in non-supplemented nests. The next best performing model included all two-way interaction terms and indicated that body condition was negatively correlated with chicks’ HPA response (effect size: -0.57 ± -1.05, -0.08).

**Figure 3.**
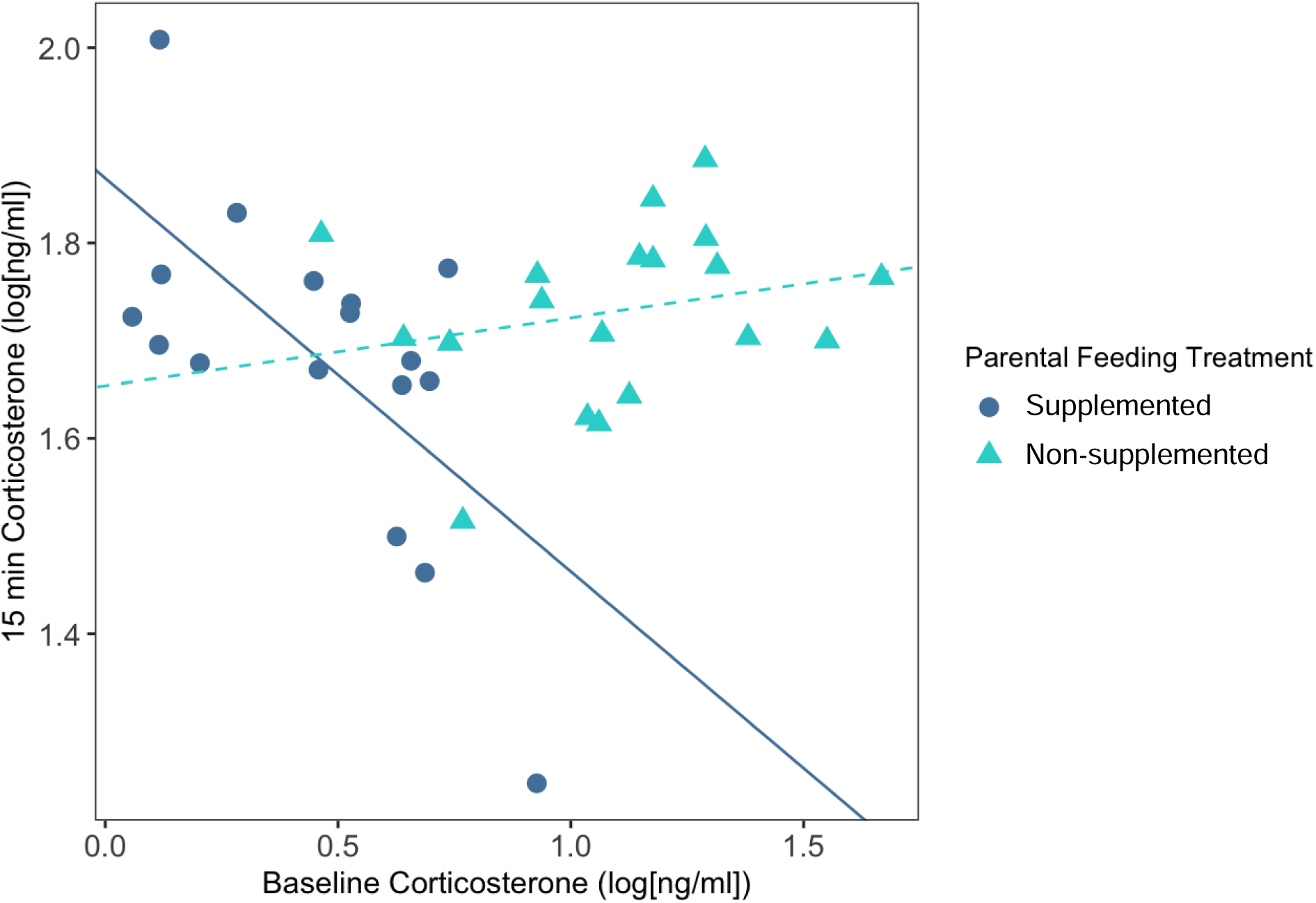
Baseline and restraint-induced corticosterone. Baseline corticosterone of 5-day old kittiwake chicks was negatively correlated with corticosterone levels after 15 min bag restraint, but only in chicks from nests where parents received supplemental feeding (“supplemented”). Chicks from supplemented nests had lower baseline corticosterone to begin with (see figure 4A). Solid line = significant, dashed = non-significant.

**Figure 4.**
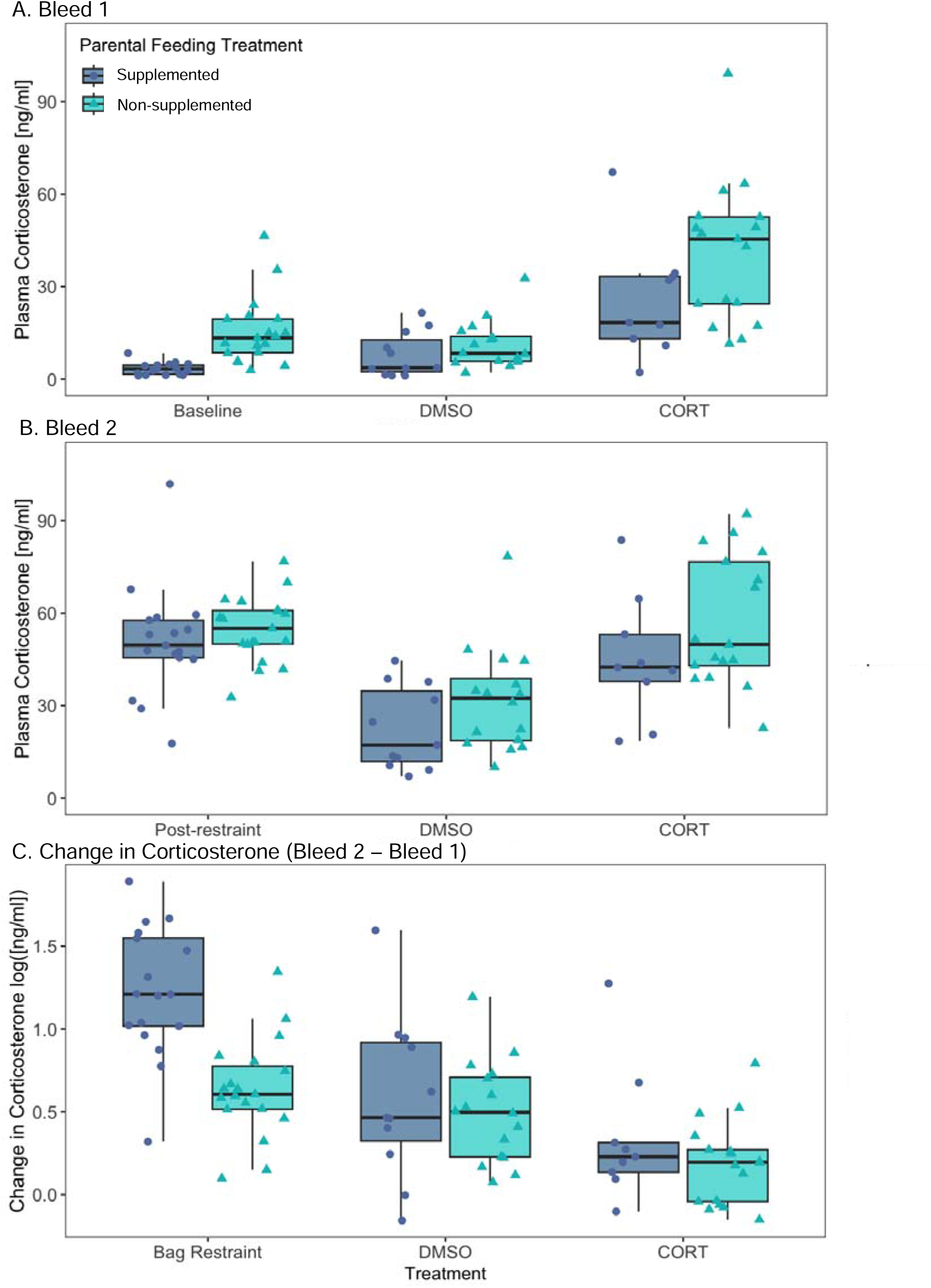
Hormone treatments and repeated blood samples. Circulating corticosterone levels in 5-day old kittiwake chicks by parental feeding status and chick treatment. A) Bleed 1 is an undisturbed baseline bleed for the control group (< 3 min) and 15 min after topical treatments for the DMSO (DMSO gel) and CORT (corticosterone dissolved in DMSO gel) groups. B) Bleed 2 is after 15 min of continuous bag restraint for the control group, and 15 min after the first bleed for the DMSO and CORT groups (treated chicks were returned to nests between bleed 1 and bleed 2). C) Change in corticosterone levels between the first and second bleed. All statistical analyses were performed using log-transformed data, but raw data are shown for panels A and B.

**Table 4.** Model selection results of variables affecting corticosterone response to 15 minutes of handling and restraint in 5-day old chicks. Baseline corticosterone is log-transformed. Treatment = food supplementation to parents (supplemented or non-supplemented). *A priori* justification is the same as in Table 3 with the addition of baseline corticosterone: a chick’s baseline physiological state affects their response to a stressor, and that response may exist in parallel or be modified by the other study variables. Since chick sex was not a significant predictor of baseline corticosterone but there were significant interactions among the other variables of interest, we included a full two-way interaction without sex in the model set. The top performing model is italicized.

| Model | K | AICc | Delta AICc | AICcWt |
| --- | --- | --- | --- | --- |
| *Treatment\*Baseline corticosterone* | 5 | -49.38 | 0 | 0.66 |
| Two-Way Interaction Model without Sex* | 8 | -47.29 | 2.09 | 0.23 |
| Body Condition | 3 | -42.28 | 7.1 | 0.02 |
| Treatment+Body Condition+Baseline corticosterone | 5 | -41.24 | 8.14 | 0.01 |
| Null (intercept only) Model | 2 | -41.23 | 8.15 | 0.01 |
| Treatment+Body Condition | 4 | -40.84 | 8.54 | 0.01 |
| Treatment+Baseline corticosterone | 4 | -40.27 | 9.11 | 0.01 |
| Treatment | 3 | -40.16 | 9.22 | 0.01 |
| Body Condition+Baseline corticosterone | 4 | -39.86 | 9.52 | 0.01 |
| Sex+Body Condition | 4 | -39.83 | 9.55 | 0.01 |
| Sex | 3 | -39.28 | 10.1 | 0 |
| Treatment*Body Condition | 5 | -38.96 | 10.41 | 0 |
| Baseline CORT | 3 | -38.87 | 10.51 | 0 |
| Treatment+Sex+Body Condition+Baseline corticosterone | 6 | -38.86 | 10.52 | 0 |
| Treatment*Body Condition+Baseline corticosterone | 6 | -38.81 | 10.57 | 0 |
| Treatment+Sex | 4 | -38.16 | 11.22 | 0 |
| Treatment+Sex+Body Condition+Baseline corticosterone | 5 | -37.67 | 11.7 | 0 |
| Treatment*Sex | 5 | -37.26 | 12.12 | 0 |
| Sex*Body Condition | 5 | -37.15 | 12.23 | 0 |
| Sex*Baseline corticosterone | 5 | -36.96 | 12.42 | 0 |
| Sex+Baseline corticosterone | 4 | -36.8 | 12.58 | 0 |
| Full Two-Way Interaction Model including Sex** | 12 | -32.27 | 17.1 | 0 |

\*Treatment+Sex+BC+BaselineCorticosterone+Treatment\*Sex+Treatment\*BC+Treatment\*BaselineCorticosterone+Sex\*BC+Sex\*BaselineCorticosterone+BC\*BaselineCorticosterone
\*\*Treatment+BC+BaselineCorticosterone+Treatment\*BC+Treatment\*BaselineCorticosterone+BC\*BaselineCorticosterone

### 3.3 Experiment 2: Exogenous corticosterone and negative feedback

#### 3.3.1 Response to exogenous corticosterone treatment

Topical application of corticosterone significantly increased plasma corticosterone (F_5, 84_ = 61.08, p < 0.0001, controlled for effect of body condition F_1, 88_ = 5.76, p = 0.018, with individual as a random effect). Post-hoc tests showed that the CORT treatment elevated circulating corticosterone compared to undisturbed baseline (p < 0.001) and to the DMSO treatment (p < 0.001), but the identical application of DMSO alone did not elevate corticosterone compared to undisturbed baseline (p = 0.998), confirming that neither DMSO nor the topical administration procedure evoked significant changes in HPA activity (Table 2; Fig 4). The topical CORT treatment increased circulating corticosterone to levels that were slightly (but significantly; p = 0.001) lower than levels secreted endogenously in response to 15 minutes of restraint (Fig 4A/B). Parental feeding treatment had a strong effect on corticosterone levels 15 min after exogenous corticosterone application (F_1,50_ = 4.56, p = 0.037, controlling for body condition F_1,50_ = 0.34, p = 0.56) -- chicks from supplemented nests had much lower circulating corticosterone 15 min after the topical hormone application than chicks from non-supplemented nests (Table 2; Fig 4). This difference between supplemented and non-supplemented groups mirrored the difference between them in the baseline group: i.e. corticosterone in non-supplemented chicks was 12 ng/mL higher than in supplemented chicks at baseline. A separate small subset of chicks bled 5 min after gel application showed no significant difference between the DMSO and the CORT treatments (t_5_ = -1.07, p = 0.33; Table 2).

#### 3.3.2 Effects of exogenous treatment on endogenous response to handling

Both the CORT and DMSO-treated chicks increased circulating corticosterone at the second bleed (30 min after topical gel application and 15 min after the first bleed) compared to their respective first bleeds (post-hoc comparison from mixed-effects ANCOVA, DMSO p < 0.001, CORT p = 0.003; Table 2; Fig 4C). Although the corticosterone in the CORT treated group remained higher at the second bleed compared to the DMSO group (p < 0.001; Fig 4B), the adrenocortical response (magnitude of change) was larger in the DMSO treatment (F_1,49_ = 8.6626, p = 0.005; Fig 4C).

In addition, DMSO and CORT treated chicks from supplemented nests had lower corticosterone at both the first (F_1,50_ = 5.17, p = 0.03) and second bleed (F_1,50_ = 4.56, p = 0.04) compared to those from non-supplemented nests (in both cases controlling for body condition, though body condition was not significant; bleed 1: F_1,50_ = 0.12, p = 0.45; bleed 2: F_1,50_ = 0.02, p = 0.56; Fig 4A/B). However, feeding treatment did not affect negative feedback. The differences in corticosterone levels between parental feeding treatments at the second bleed were of the same magnitude as the difference between feeding treatments at the 1st bleed (15 min after gel application; F_1,49_ = 1.2688, p = 0.26). Consequently, there were no differences between feeding treatments in the magnitude of change in corticosterone within CORT or DMSO treatments (bleed 2 corticosterone - bleed 1 corticosterone; Fig 4C). Finally, the increase in endogenous corticosterone after the brief disturbance of handling followed by return to the nest for ∼ 15 min (delta corticosterone for the DMSO treated group) was much lower than the endogenous increase in corticosterone after 15 min of continuous bag restraint (F_2.82_ = 21.29, p < 0.0001; Fig 4C).

### 3.4 Experiment 3. Clearance of exogenous corticosterone

In non-supplemented nests in 2022, circulating corticosterone in DMSO-treated chicks bled for the first time 15 or 60 min after treatment did not differ from undisturbed baseline corticosterone (Tukey Contrasts, p = 1 and 0.63 respectively; Fig 5, Table 2). Corticosterone in the CORT treatment was higher than baseline and DMSO at both 15 (p = 0.01 and 0.007) and 60 min post-treatment (p = 0.02 and <0.001) but was not different between the 15- and 60-min CORT groups (p = 0.97; Fig 5).

**Figure 5.**
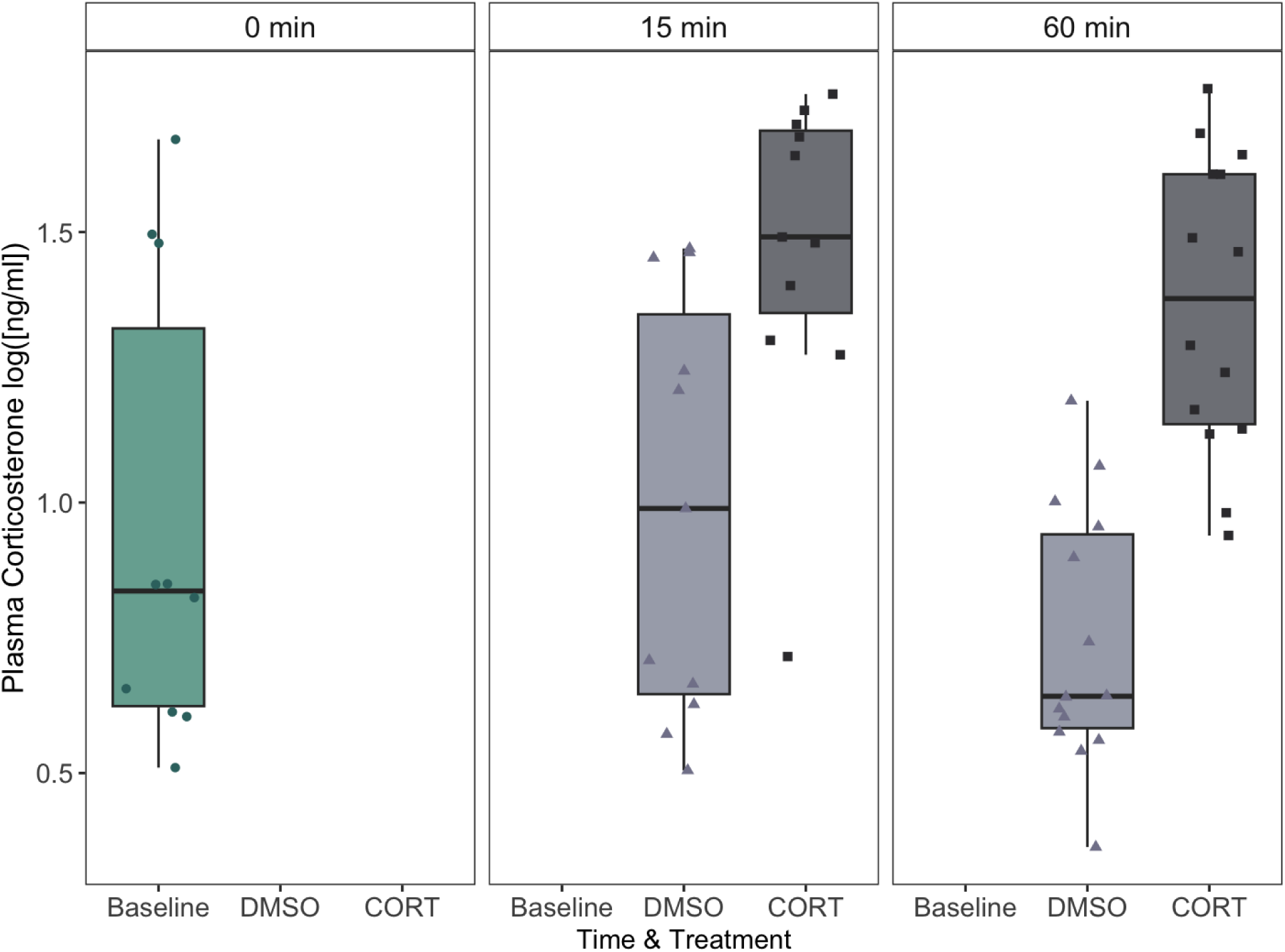
Time course for topical corticosterone. Corticosterone levels in 5-day old chicks from nests without parental food supplementation (non-supplemented) at undisturbed baseline (< 3 min), 15 and 60 minutes after application of topical gel treatments. No chick was bled more than once. Statistical analyses were performed using log-transformed data but raw data are shown here.

## 4. Discussion

In this study, we demonstrate that kittiwake chicks do not exhibit a post-hatch HPA-axis hypo-responsive period. Instead, they have HPA axes that a) can rapidly secrete a large amount of corticosterone and b) are sensitive to environmental and physiological context. At five days old, just 10-15% of the way through the nestling stage, chicks mounted corticosterone responses proportional to the intensity of the challenge, with levels 15 min after a moderate challenge (brief handling to take a blood sample followed by return to the nest) that were higher than the undisturbed baseline, but lower than levels evoked by a more intense challenge (15 min of continuous restraint). Chick corticosterone levels after the 15 min restraint challenge were not only much higher than their baseline levels, but higher than any of the previously reported maximum values for kittiwake chicks with the exception of much older, severely food-restricted captive chicks (though all previous studies that measured restraint-induced levels used chicks older than 5 days; Table 1). Restraint-induced levels of corticosterone in chicks were comparable to (with some individuals higher than) maximal levels secreted by adult kittiwakes at this colony during 50 min of restraint (12 - 63 ng/mL; Kitaysky et al., 2010). In addition, nutritional status of parents interacted with interannual variability and body condition to affect baseline HPA axis activity, though neither chick sex nor number of siblings did. Our study design minimized potential sources of variation in HPA activity by using kittiwake chicks of known age, hatching order and sex, and sampling within a narrow hatch date range (< 2 weeks) and time of day (< 3 hours).

### 4.1 Food availability to parents and year effects

Despite the fact that 5-day old chicks are very early in development and parents should be able to meet their relatively low food needs at this stage, there were strong effects of year and parental supplemental feeding treatment on chick HPA activity. Baseline corticosterone was significantly lower in chicks from food supplemented nests in one of two years. An obvious explanation for this difference might be that the current nutritional status of parents is transmitted to chicks via differences in provisioning, which affects chick nutritional status and thereby their HPA activity. However, two lines of evidence indicate that this was a contributing factor but not the sole driver. First, parental food treatment had a strong effect in 2021 even when chick body condition was controlled for statistically. Second, there were no significant differences in chick body condition between years or feeding treatments, yet their baseline circulating corticosterone levels differed between years. In fact, although climate indices and measures of reproductive success indicated that 2021 and 2022 were both years of intermediate reproductive success for kittiwakes at this site, 2022 was a better year than 2021 (S. Hatch, unpublished data). Yet baseline corticosterone was higher in 2022 than in 2021, and in 2022 chicks from supplemented nests had corticosterone levels that were similar to non-supplemented. Though reproductive success at the end of the season is temporally distant from conditions at 5 days post hatch, the mismatch between annual breeding success and young chick endocrine physiology suggest that the differences in chick baseline corticosterone are affected by food availability but also modulated by additional factors that we were unable to measure in this study.

There are several potential modulators of the relationship between chick corticosterone and current food availability. First, it could be a result of prenatal effects. Supplemental feeding at experimental nests began before egg-laying, potentially interacting with early season or even overwinter environmental conditions in each year to influence egg composition (Benowitz-Fredericks et al., 2013). Variation in egg composition can affect chick HPA activity (Ahmed et al., 2014; Love and Williams, 2008; Schwabl and Partecke, 2020; Sockman and Schwabl 2001) and may be a mechanism for intergenerational transmission of information that prepares chicks for the environment into which they are born (reviewed in Engqvist and Reinhold, 2016). In addition, adults show high nest site fidelity at this colony, thus most parents at supplemented and non-supplemented nests have likely experienced the same feeding treatments in past years. This combination of early indicators of natural food availability during the pre-lay period (Whelan et al., 2021) and anticipated changes in food availability during chick-rearing based on prior experience may shape investment in reproduction (Shultz et al., 2009), and thus the programming of chicks’ HPA axes. Second, the differences in HPA activity could be driven by post-natal behavioral cues from parents or siblings. While chick body condition indicated that parents seemed to be meeting the nutritional needs of their chicks at this early age regardless of the supplemental food treatment, differences in competition from siblings or responsiveness of parents to begging may impact corticosterone secretion (Braasch et al., 2014; Dreiss et al., 2016). Finally, other aspects of interannual environmental variability could modulate the relationship between food availability and corticosterone by imposing additional energetic challenges. For example, weather can affect chick corticosterone (Crino et al., 2020; De Bruijn and Romero, 2018), as can ectoparasites (Dudek et al., 2021; Kitaysky, 2001; Quillfeldt et al., 2004).

### 4.2 Body condition

Control chicks (from non-supplemented nests) in higher body condition had lower baseline corticosterone levels, and body condition was generally negatively correlated with restraint-induced corticosterone levels. While negative relationships between body condition or body mass and corticosterone have been frequently reported in birds, including adult black legged kittiwakes (Kitaysky et al. 1999) and red-legged kittiwake chicks (Kitaysky et al. 2001), a phylogenetically controlled meta-analysis found no consistent relationship between baseline corticosterone and body condition in seabirds (Sorenson et al., 2017). In adult kittiwakes, low body condition can be associated with either low or high levels of corticosterone (Schultner et al., 2013) - the first because animals with a dependable food supply and low corticosterone can afford to avoid the costs of carrying extra weight (Rogers, 2015), and the second because severe food shortages elevate corticosterone, which mobilize and deplete reserves. Growing chicks are under different selection pressures than adults and may be less likely to benefit from maintaining a “lean” phenotype, though the relationships between growth and fitness in kittiwakes are complex and context-dependent (Vincenzi et al., 2013; Vincenzi and Mangel, 2013). Indeed, in this experiment the relationship between body condition and baseline corticosterone was eliminated by supplemental feeding. While some models predict that the relationships between glucocorticoids and phenotype will be obscured by high resource availability (Breuner and Berk, 2019), there were no differences in body condition between treatments in the young chicks included in this study. This indicates that at least at this age, factors in addition to internal indicators of energy reserves may modify chick HPA activity. For example, other aspects of interannual variability, such as weather, may be buffered to different extents by parents in the different supplemental feeding treatments. The negative relationship with response to restraint indicates that chicks in better condition may be able to buffer themselves from potential costs of elevated glucocorticoids exposure by downregulating their sensitivity to challenges. However, it is also worth noting that food intake can change tissue composition of chicks including increased water retention (Kitaysky et al., 1999) and, potentially, differences in rate of yolk absorption (Noy and Sklan, 2001), such that measures of body condition in young chicks may not always provide an accurate proxy of nutritional status.

### 4.3 Genetic sex

There are relatively few studies examining sex differences in chick HPA activity. While sexual dimorphism in other traits might be hypothesized to predict sexual dimorphism in physiology, there is variation among species in whether chick HPA axis activity varies by sex, with some showing sex differences in HPA responsiveness before dimorphism emerges (e.g. Lynn et al. 2022) and others showing no differences despite sexual dimorphism (e.g. Sockman and Schwabl 2001). Though young kittiwake chicks are initially sexually monomorphic, males grow faster and are generally considered more costly for parents to rear (Merkling et al., 2012; Young et al., 2017). While the dimorphism manifests when chicks are older than 5 days (Merkling et al. 2012), the physiological substrates for these differences may be established earlier. One study measured baseline corticosterone in 5-day old kittiwakes and, similar to this study, found no effect of sex. However, none of the additional aspects of HPA-function that we measured in this study (response to restraint, negative feedback) varied by sex either. Whether a sex difference manifests later in development as dimorphism increases, or whether the effects of elevated corticosterone on physiology and behavior differ by sex even when HPA activity does not, remains to be determined.

### 4.4 Sources of variation in restraint-induced corticosterone

The magnitude of the corticosterone response to restraint in chicks has been proposed to reflect consistent phenotypic variation in adrenocortical responsiveness with fitness implications, predicting subsequent recruitment and survival in at least one species (Blas et al. 2007). What drives variation in restraint-induced corticosterone levels in chicks? Though we did not measure it here, HPA axis activity has a heritable component in many birds, with higher heritability of restraint induced levels than baseline levels (Beziers et al. 2019, Jenkins et al. 2014). While body condition may also play a role (see section 4.2), the strongest evidence in this study was for a negative relationship between baseline corticosterone and corticosterone after restraint, but only among chicks in the food-supplemented treatment. Chicks from supplemented nests secreted more corticosterone to reach similar levels as their counterparts from the non-supplemented nests after 15 min of restraint, despite starting at much lower baseline levels. While these results suggest that lower baseline corticosterone levels in chicks from supplemented nests was not a consequence of a difference in adrenal capacity, they do not identify sources of interindividual variation within treatments.

Variation in adrenal capacity may be explained in part by the “facilitation” phenomenon, in which restraint-induced corticosterone levels reflect adrenal capacity as shaped by HPA activity in the recent past (Akana et al., 1992; Dallman et al., 2004, 1992; Kitaysky et al. 2007). In a study of adult kittiwakes, maximum restraint-induced corticosterone was correlated with food availability two weeks prior to sampling (Kitaysky et al., 2010). Though the adrenal capacity of very young chicks has had limited opportunity to be shaped by post-hatch experience, the HPA axis of avian embryos is active and capable of responding to stimuli (Jenkins and Porter, 2004) and its activity in chicks may be shaped by both prenatal and postnatal exposure to corticosterone (Benowitz-Fredericks et al. 2015; Marasco et al., 2016), as well as environmental context (Groothuis et al., 2020). Thus, it is possible that prenatal experience plays a role (see section 4.1) and the HPA responsiveness of chicks to restraint is determined by a combination of ontogenetic changes and embryonic/early post-hatch exposures to external cues.

### 4.5 Effects of exogenous glucocorticoids

We used topical corticosterone administration to evaluate rapid uptake, clearance, and negative feedback of glucocorticoids. The ability to clear circulating glucocorticoids via a combination of negative feedback and metabolism is an important aspect of HPA function that likely has fitness consequences, as it allows animals to buffer themselves from costs of extended glucocorticoid exposure (Lattin and Kelly, 2020; Romero, 2004; Romero et al., 2009; Taff et al., 2018). Though the effects were not apparent after 5 minutes, topical corticosterone treatment elevated circulating corticosterone within a physiologically relevant range after 15 min. This elevation persisted for at least 60 min. In contrast, administration of the vehicle control (DMSO alone) did not change corticosterone levels, compared to undisturbed baseline levels, at any point.

We did not find evidence that chicks from supplemented and non-supplemented nests processed exogenous corticosterone differently. The difference between parental feeding treatments in levels of corticosterone 15 min after administration of exogenous corticosterone was similar to the difference between them at baseline. However, the second bleed for the DMSO and CORT treated chicks provided evidence that a) exogenous corticosterone evoked rapid negative feedback on endogenous production (the magnitude of the handling-induced increase was higher in the DMSO control treatment compared to the CORT treatment) and b) the levels of negative feedback did not differ by parental feeding treatment (the magnitude of difference in corticosterone between CORT treated chicks in supplemented and non-supplemented nests at the 2nd bleed were the same magnitude of difference as in the first bleed). Thus, while endogenous secretion of corticosterone in 5-day old chicks is sensitive to environmental context, rapid negative feedback and/or the ability to clear acute exogenous elevations in corticosterone are not. Though altered sensitivity to negative feedback has been proposed as a potential mechanism to explain ontogenetic changes in HPA activity in chicks, this hypothesis was not supported in white storks (TorresLMedina et al., 2019). Similarly, our study suggests that it does not explain variation in HPA activity between young kittiwake chicks from supplemented and non-supplemented nests, though age-related changes have not yet been tested.

### 4.6 Conclusion

The developmental trajectory of avian HPA axes has been hypothesized to be driven by a species’ position on the altricial-precocial continuum (Blas et al., 2006), latitude (Jones et al., 2021), or by other aspects of life-history including provisioning strategies, but no consistent patterns have emerged (Sagar et al., 2019). The HPA axes of black-legged kittiwake chicks are sensitive to internal and external cues and are capable of mounting a large glucocorticoid response to an acute challenge (restraint) during a developmental window of intense social interaction with both siblings and parents. While this is consistent with the hypothesis that glucocorticoids are important for species with intra-brood competition, the ability of young chicks to mount an adult-like HPA response to restraint has been found in a number of other seabird species that have one-egg clutches, including three species of petrels (Adams et al., 2008; Angelier et al., 2022; Sagar et al., 2019), three species of alcids (Benowitz-Fredericks et al., 2015; Kitaysky et al., 2005; Will et al., 2015), and a congener of the black-legged kittiwakes studied here: the red-legged kittiwake (Kitaysky et al., 2001; though red-legged kittiwakes historically had larger clutch sizes). Thus, sibling competition cannot be the sole selective pressure on the development of HPA axis activity. A phylogenetically controlled meta-analysis characterizing the ontogeny of HPA activity across species with varied ecologies and life-histories would be valuable for understanding the pressures and constraints on HPA axis development in birds. However, for species like kittiwakes with very early activation of the axis, the next steps are to understand why young chicks are capable of such robust responses to acute challenges by characterizing the rapid metabolic and behavioral effects of elevated corticosterone in chicks.

## Acknowledgements

We are grateful for help with the following: establishment, maintenance, and long-term funding/logistics for the tower colony: S. Hatch and M. Hatch; field logistics and assistance - F. Tremblay and E.C. Kelly. Field crews: 2021 - K. Bacile, A. Colligan, J. Dunoyer, E. Fernandes-McDade, J. Morales Valenzuela, S. Morrow. 2022 - G. Dennis, H. Gee, L. Jackson, D. Jadhon, L. Marcouillier, S. Solasz, A. Turmaine; impeccable hormone assays - E. Kitaiskaia; genetic sexing - A. Jackson, A. Le, S. Lin, and S. Walsh. We also thank two anonymous reviewers for their thoughtful feedback. Financial support came from a Dean’s Fellowship from Bucknell University (ZMBF), a Gates Millennium Scholarship and an American Indigenous Business Leaders internship (SNP), Gulf Watch Alaska and Alaska Ocean Observing System (SW/Institute for Seabird Research and Conservation), and the Institute of Arctic Biology, University of Alaska Fairbanks (ASK and AW). All research was conducted under permits from the U.S. Fish and Wildlife Service (#MB33779D-1), Alaska Department of Fish and Game (#21-089; #22-080) and with approval from McGill University’s Animal Use Committee.

